# Enhancing Protein Binding Site Residue Prediction with Graph Neural Networks: Impacts of Cutoff Distance and Feature Selection

**DOI:** 10.1101/2025.08.25.672254

**Authors:** Serena H. Chen, Massimiliano Lupo Pasini, Cory D. Hauck

## Abstract

Identifying protein binding site residues is critical for understanding molecular interactions but challenging due to incomplete knowledge of features that determine binding. The increasing volume and accessibility of protein sequences and structures presents an opportunity to systematically decipher their intricate relationships with binding. Here, we investigate the effect of cutoff distance between residues and the importance of sequence-based and structure-based features on binding in four eukaryotic model organisms with graph neural networks. Our results indicate that sparse graphs generated by an 8 Å cutoff are most effective for binding site residue prediction, and that increasing cutoff distance introduces noise from non-binding site residues, impairing model performance. Yet, using hybrid cutoffs that enrich connectivity between predicted binding site residues has the potential for enhancing performance. In addition, sequence-based features learned by a pre-trained protein language model carry substantial information to binding. In contrast, structure-based features derived from protein backbone geometry are inadequate due to overprediction of binding site residues. These findings are consistent for the proteins across the four organisms, providing evidence for evolutionary conservation of binding site residues.

This manuscript has been authored by UT-Battelle, LLC under Contract No. DE-AC05-00OR22725 with the U.S. Department of Energy. The United States Government retains and the publisher, by accepting the article for publication, acknowledges that the United States Government retains a non-exclusive, paid-up, irrevocable, worldwide license to publish or reproduce the published form of this manuscript, or allow others to do so, for United States Government purposes. The Department of Energy will provide public access to these results of federally sponsored research in accordance with the DOE Public Access Plan (http://energy.gov/downloads/doe-public-access-plan).

## Main

Binding is a fundamental mechanism of protein function. This process involves the interaction of a protein with other molecules including nucleic acids, carbohydrates, lipids, metals, and other proteins. Binding interactions are driven by a protein’s three-dimensional (3D) structure, which is largely determined by the residues in its primary sequence. Identifying which residues are responsible for binding is an essential step to understand molecular mechanisms and to create or modulate binding interactions.

Recently, graph neural networks (GNNs) have shown promise in predicting protein binding site residues [1–5]. Graphs are a natural data structure for representing protein 3D structures, where nodes represent residues and edges are defined based on the spatial relationships between residues. Message passing algorithms that are core to the training of GNNs provide a logical approach for capturing pairwise residue interactions. When representing proteins as graphs for GNN training, at least two important issues arise: (i) the choice of cutoff distance that defines local residue neighborhoods for message passing and (ii) the choice of features that are used for inputs.

Cutoff distances are often applied when computing nonbonded interactions in molecular dynamics (MD) simulations of proteins to improve computational efficiency. Nonbonded interactions consist of van der Waals (vdW) and electrostatic forces which decay with increasing distance. With cutoff distances, any interaction between protein atoms separated beyond a defined cutoff is either neglected or approximated. These approximations also apply to graphs when constructing edges. The effect of cutoff distances have been evaluated on protein structures in MD simulations [6] and on protein graphs for select applications [7–10], yet their effects on protein graphs for binding site residue prediction has not been systematically studied.

Features are commonly extracted from protein sequences and structures. Protein sequences change over time due to random mutation and natural selection. Yet, functionally important residues, such as binding site residues, are often preserved for organisms’ fitness, creating signatures that may be extracted from protein sequences [11]. Similarly, protein structures are also conserved to maintain specific functions [12]. Even non-homologous proteins may have independently developed similar structures and functions [13]. As a result, protein sequences and structures carry informative features for binding site residue prediction.

Commonly used sequence-based and structure-based features are either handcrafted by domain knowledge or learned by machine learning models. Handcrafted features rely on domain knowledge from geometric, biological, physical, and chemical features of experimentally characterized binding site residues [14–16]. Learned features rely on machine learning models to extract relevant features from data [17, 18]. For example, protein language models using long short-term memory or Transformer architectures have been used to extract residue-level features from protein sequences [5, 19]. Despite numerous feature options, it is unclear what sequence-based and structure-based features are important for binding site residue prediction.

Existing GNNs for protein binding site residue prediction lack a consensus in the choice of cutoff distance and features. GraphBind [1] focuses on predicting protein-nucleic acid binding site residues using graphs constructed with a 10 Å-cutoff and a set of handcrafted features derived from evolutionary conservation, secondary structure, residue position, and residue’s atomic features, such as mass, charge, vdW radius and other physicochemical features. Although two other predefined cutoff distances, 5 Å and 13 Å, were evaluated, the 10 Å-cutoff was found to be the most favorable for nucleic acid binding. Another GNN, GraphPPIS [2], predicts protein-protein binding site residues using features from evolutionary conservation and secondary structure. GraphPPIS treated the cutoff distance as a hyperparameter and found that a 14 Å-cutoff was the best performing. GeoNet [4] and GPsite [5] target protein-ligand binding site residue prediction. Both selected an arbitrary cutoff distance, with GeoNet using a 10 Å-cutoff and GPsite using a 15 Å-cutoff. It is likely that the different cutoff distances result from the different message passing algorithms employed. However, it raises a question whether, given the same message passing algorithm, there exists an optimal cutoff distance regardless the type of binding interactions. The majority of these GNNs use handcrafted features which are often inter-correlated. For instance, evolutionary conservation features are expected to correlate with secondary structure features given the conservation patterns of certain secondary structure elements [12]. Moreover, atomic charge and vdW radius are directly related to the residue’s solvent accessible surface area. Feature selection requires careful evaluation since the most orthogonal features contribute the most information to GNN’s prediction.

In this work, we systematically investigate cutoff distance and feature importance for binding site residue prediction with GNNs (**Fig. 1**). We find that an 8 Å cutoff is most effective and that increasing the cutoff distance introduces noise from non-binding residues and impairs model learning. We discover that using hybrid cutoff distances to increase message passing between predicted binding site residues has the potential for enhancing model performance. We also find that structure-based features derived from protein backbone geometry encode some binding information, but using such features alone generates numerous false positives. On the other hand, a GNN using sequence-based features learned from a pre-trained protein language model achieves similar performances as a GNN using both sequence-based and structure-based features. This result suggests that the learned sequence-based features not only encode some level of backbone geometry but also carry important information not contained in the structure-based features for binding. We observe that GNNs trained on human proteins are generalizable to mouse, rat, and yeast proteins, providing evidence for evolutionary conservation of binding site residues.

**Fig. 1:**
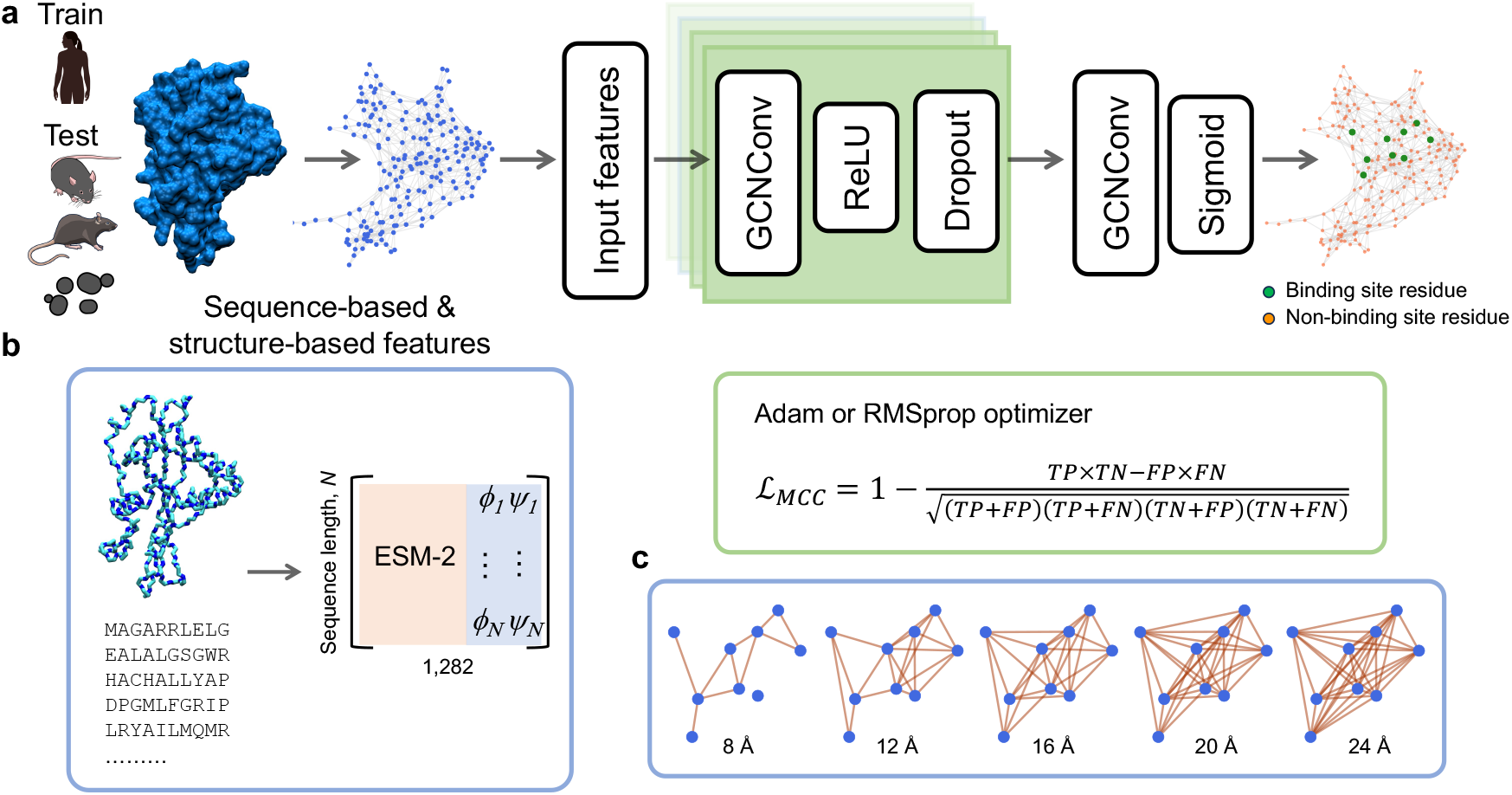
Investigating cutoff distance and feature importance for protein binding site residue prediction with graph neural networks. **a** Overview of the workflow for protein binding site residue prediction. Proteins are converted into graphs. Sequence-based and structure-based features are the input of the graph neural networks (GNNs). The GNNs use one to four hidden layers with each layer consisting of a GCNConv graph convolutional operator with ReLU activation function, followed by dropout regularization during training. The output layer consists of a GCN-Conv graph convolutional operator with the sigmoid activation function to generate a probability of being a binding site residue for each residue of the proteins. The GNNs are trained on the human proteins using the Adam or RMSprop optimizer to minimize Matthews Correlation Coefficient loss function (ℒ_*MCC*_) defined by true positives (TP), true negatives (TN), false positives (FP), and false negatives (FN). The GNNs are then tested on the mouse, rat, and yeast proteins. Illustrations of human, mouse, and rat are obtained from NIAID Visual & Medical Arts entries 227 [20], 281 [21], and 54 [22], respectively. **b** Each node is characterized by sequence-based features computed from the final hidden state of the pre-trained protein language model ESM-2 and by structure-based features using the backbone dihedral angles *ϕ* and *ψ*. **c** Graphs of the protein from **a** showing the increasing number of edges between binding site residues at cutoff distances ranging from 8 Å to 24 Å at an interval of 4 Å.

## Results

### Dataset preparation and graph representation for protein binding site residue prediction

To study binding in relation to protein sequences and structures, we first construct datasets by combining residue-level binding annotations in the Universal Protein Resource Knowledgebase (UniProtKB) [23] with residue types and atomic coordinates in AlphaFold Protein Structure Database (AlphaFold DB) [24,25]. UniProtKB provides residue-level binding annotations of protein sequences for four model organisms: *Homo sapiens* (human), *Mus musculus* (mouse), *Rattus norvegicus* (rat), and *Saccharomyces cerevisiae* (yeast). These annotations are based on expert curation of experimental data in the literature and cross references of protein family and domain annotations from curated databases [26, 27]. A residue is annotated as binding site residue if it interacts with any chemical entities, including metals, cofactors, activators, inhibitors, regulators, and enzyme substrates and products [23]. We select proteins with at least one annotated binding site residue and collect their Protein Data Bank (PDB) files from AlphaFold DB. AlphaFold DB provides protein 3D structures predicted by AlphaFold2 [24] in PDB format for protein sequences from UniProtKB. The PDB files include residue types and atomic coordinates that we use to compute features for binding site residue prediction. By pairing protein sequences and protein structures from AlphaFold DB with the corresponding binding annotations from UniProtKB, we construct a dataset for proteins with at least one binding site residue for each of the four model organisms. We label annotated residues as 1 for binding site residues and 0 for non-binding site residues. In all four datasets, binding site residues are extremely rare, consisting of between 1.6% and 1.9% of the total protein residues (**Table 1**). These low percentages indicate that the datasets are highly imbalanced.

**Table 1:**
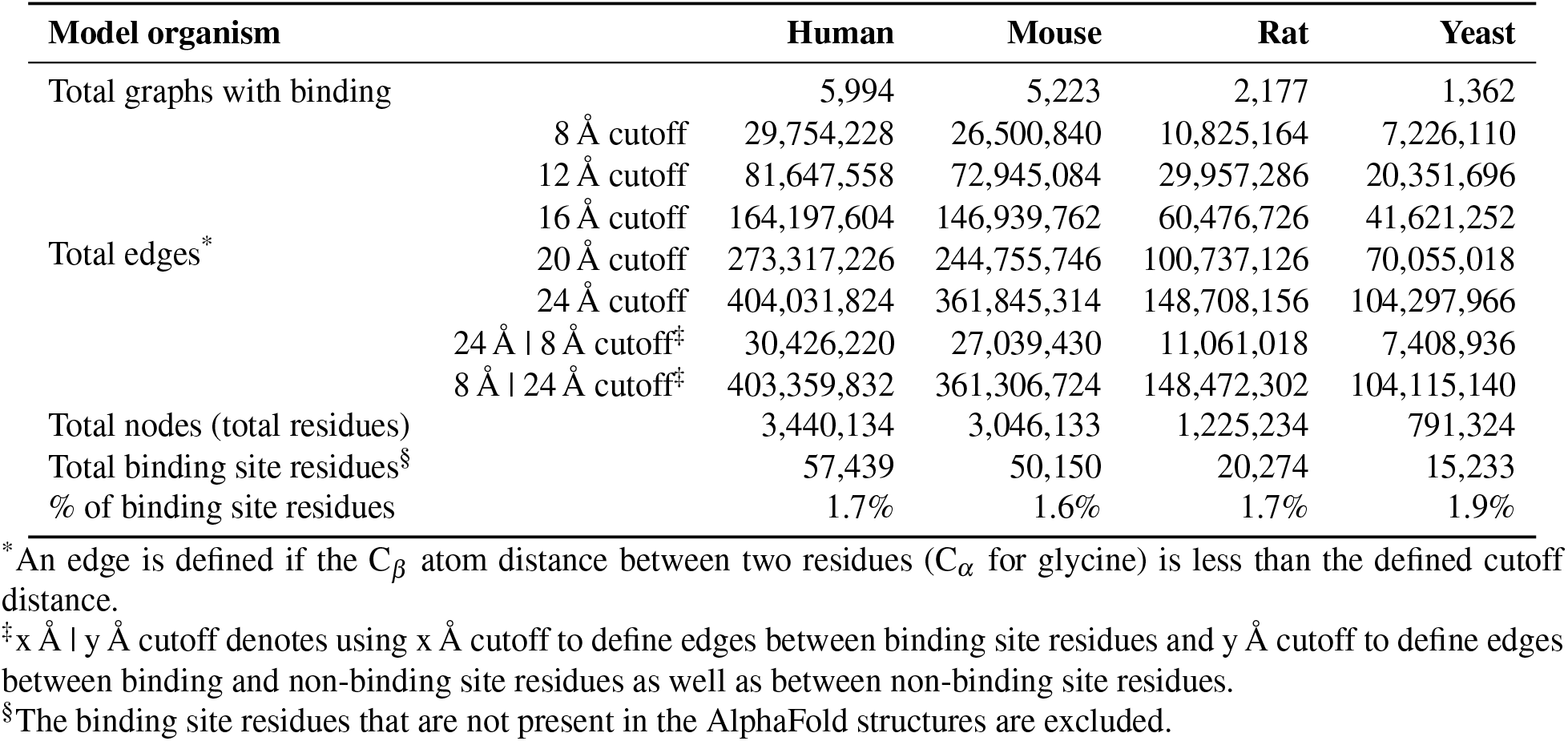
Summary of the protein graphs in the four datasets.

We convert the proteins in each of the four datasets into graphs (see Methods for details). Graph representations are useful for analyzing protein structures since the complex spatial relationships and interactions between protein residues can be efficiently captured using graph edges and nodes. We use graph nodes to represent protein residues and undirected edges to represent pairwise residue interactions. An edge is assigned between two residues if the distance between their C_*b*_ atoms (C_*a*_ atom for glycine) is less than an empirical cutoff distance. There are various approaches attempting to avoid over connectivity of graphs, such as the k-nearest neighbors algorithm and the spherical Voronoi diagram algorithm, yet distance-based methods are physically informative and remain widely used for edges of protein graphs [28]. As intermolecular forces decrease rapidly with increasing distance, we consider cutoffs from 8 Å, 12 Å, 16 Å, 20 Å, and up to 24 Å (**Fig. 1c**). 8 Å and 12 Å are commonly used cutoffs for nonbonded interaction calculations in force field development and molecular dynamics simulation of proteins [29–32]. 24 Å is the upper bound given that the average distance between pairs of binding site residues levels off by 24 Å in all datasets (**Fig. S1**). As expected, the number of edges increases with increasing cutoff distance (**Table 1**), scaling up to a polynomial of degree 3.

We associate to each node a set of features extracted from the protein sequence and protein structure (**Fig. 1b**). For sequence-based features, we represent each residue by the final hidden state, a 1,280-dimensional vector, of a pre-trained Evolutionary Scale Modeling ESM-2 Transformer [33]. ESM models are proven to be powerful protein language models that extract informative features from diverse sequences across evolution [34]. These learned features encode broad biological properties from biochemical properties of residues, protein homology, to protein secondary and tertiary structures [34]. Here, we explore the importance of these features for binding site residue prediction.

Given that protein 3D structure plays an important role in binding, we seek to determine structure-based features that complement the sequence-based features for binding site residue prediction. For these features, we use the Cartesian coordinates of each residue’s backbone atoms to compute residue backbone dihedral *ϕ* and *ψ* angles. Many recent protein structure prediction methods have demonstrated remarkable success in decoding protein 3D structures from sequences alone [24, 33, 35]. Therefore, we suspect that the sequence-based features may implicitly carry some structural information. Dihedral angles *ϕ* and *ψ* are the most simple structural information and as the angles are defined by protein backbone atoms, potential sequence effects contributed by the sidechain atoms are minimized.

Sequence-based and structure-based features of the protein graphs are the input of GNNs which process and learn the relationships between nodes, mapping their relationships to their labels and returning as output the probability that each node is a binding site residue (see Methods for details). We use the graph convolutional networks based on GCNConv [36] to operate on the node features, passing the information of the computed sequence-based and structure-based features between the neighboring nodes defined by the edges. The nodes receive this information from each of their neighbors and consolidate the information before passing it back. During this iterative process, each node learns from its local graph structure, and this aggregated information is converted by a sigmoid function into a probability for being a binding site residue as the output for the node (**Fig. 1a**). We optimize the hyperparameters and train the parameters of GNNs using the human dataset. We then test the GNNs on the other three organism datasets to explore whether the models can be generalized to non-human proteins. Since the datasets are overly imbalanced, the typical accuracy loss is uninformative. Indeed, trivially classifying all nodes as non-binding site residues can achieve an accuracy close to one. Alternatively, we employ Matthews Correlation Coefficient (MCC) loss function, which is a more informative metric for binary classification tasks with imbalanced data [37–39].

### Distance-based edges supports sparse graphs for binding site residue prediction

To determine the contribution of local residue neighborhoods on binding site prediction, we evaluate five increasing cutoff distances to construct graphs with different numbers of edges for the same protein. We first investigate the composition of edges for each cutoff distance. An edge can be between two binding site residues (B-B), between a binding and a non-binding site residue (B-NB), or between two non-binding site residues (NB-NB). Given the low percentage of binding site residues, B-B and B-NB edges constitute < 5% of the edges in each dataset regardless of the cutoff distance. With increasing cutoff distance, the percentage of B-NB edges increases while the percentage of B-B edges decreases (**Fig. 2a**).

**Fig. 2:**
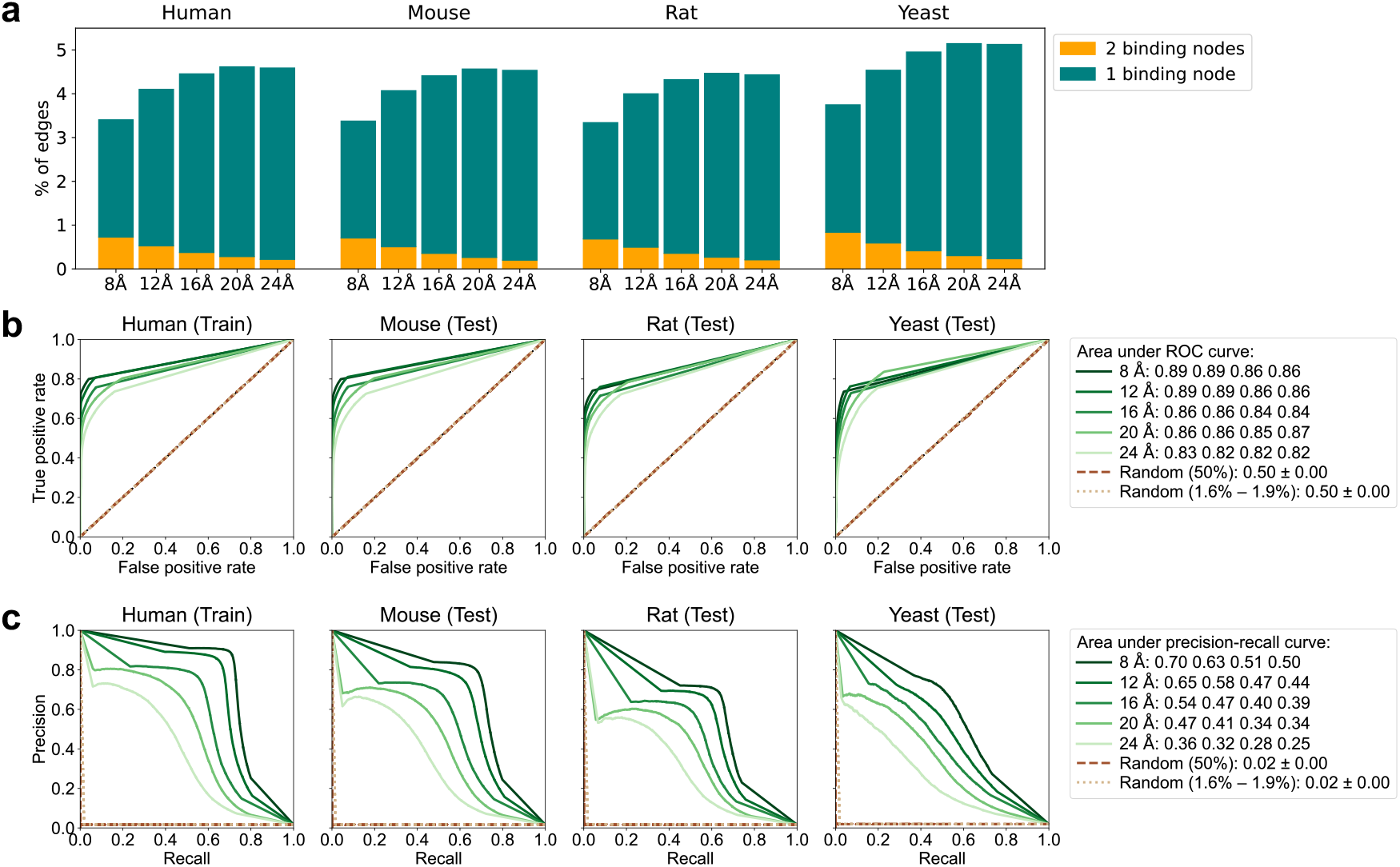
Performance of graph neural networks decreases with increasing cutoff distances. **a** Percentage of the edges between two binding site residues (orange) and between a binding and a non-binding site residues (teal) from each cutoff distance of the protein graphs in each dataset. **b** Receiver operating characteristic (ROC) curves and **c** precision-recall curves for graph neural networks (GNNs) after the last training epoch. The GNNs are trained on the human dataset and tested on the mouse, rat, and yeast datasets. All protein graphs are constructed with the cutoff distance of 8 Å, 12 Å, 16 Å, 20 Å, and 24 Å throughout the proteins, and their ROC and precision-recall curves are shown as a series of green shades from dark to light. The curves for two random guessing strategies, “Random (50%)” and “Random (1.6% – 1.9%)” are shown as brown dashed lines and tan dotted lines, respectively. The areas under these curves for the four datasets are included in the legends.

As the cutoff distance increases, more edges between residues that are further apart are included, leading to denser graphs where meaningful edges may be obscured. To explore how the cutoff distance affects GNN’s performance, for each cutoff distance, we optimize hyperparameters and parameters of a GNN on the human protein graphs and test the GNN on the mouse, rat, and yeast protein graphs. We observe consistently higher MCC loss for graphs of larger cutoff distances both on the training dataset and on the three testing datasets (**Fig. S2**). The higher loss suggests that using a larger cutoff distance, and thereby adding disproportionately more B-NB and NB-NB edges, reduces the GNN’s ability to learn and generalize binding site residues in graphs.

We also compute the receiver operating characteristic (ROC) and precision-recall curves for each of the five GNNs after the last training epoch. The area under the ROC curve measures the GNN’s ability to distinguish between binding and non-binding site residues. The area under the ROC curve is overall high, with at least a value of 0.82 (**Fig. 2b**). Among the five cutoff distances, the area under the ROC curve is greatest when using 8 Å or 12 Å for the human, mouse, and rat protein graphs. For the yeast protein graphs, the area under the ROC curve is greatest when using the 20 Å cutoff, but then decreases when the cutoff is increased to 24 Å. The variation of the area between the five cutoff distances is small in all four datasets, with a standard deviation less than 0.03.

Compared with the area under the ROC curve, the area under the precision-recall curve measures the GNN’s ability to achieve both high precision and high recall on predicting binding site residues. This metric is useful for evaluating classifiers on imbalanced datasets, where the positive class is rare. We observe a clear decreasing trend in the area under the precision-recall curve with increasing cutoff distances for all four datasets (**Fig. 2c**). Using the 8 Å cutoff, the areas are 0.70, 0.63, 0.51, and 0.50 for the human, mouse, rat, and yeast protein graphs, respectively, and these areas decrease to 0.36, 0.32, 0.28, and 0.25 when the cutoff increases to 24 Å. These results demonstrate that additional edges derived from larger cutoff distances are ineffective for the GNN to learn and predict binding site residues.

To determine if these GNNs perform better than random classification, we employ two random guessing strategies as baselines. For one strategy, named “random (50%)”, each node has a 50% probability of being a binding site residue. Yet, as we already know that binding site residues are extremely rare, all nodes do not have an equal chance of being a binding site residue. Alternatively, for the other strategy, named “random (1.6% – 1.9%)”, we apply the percent binding site residues in each dataset, randomly selecting 1.7%, 1.6%, 1.7%, and 1.9% of the nodes in each graph as binding site residues for the human, mouse, rat, and yeast datasets, respectively. The areas under the ROC and precision-recall curves using either strategy are significantly lower than any of the GNNs at 0.50 and 0.02, respectively, suggesting that the GNNs, even for the one with the lowest performance, learn from the graphs and make more informed predictions than random guessing.

### Connectivity between binding site residues is important for binding site residue prediction

To investigate the contribution of different type of edges on binding site residue prediction, we vary the composition of edges by using two cutoff distances to construct hybrid graphs for the same protein. We use a 24 Å cutoff for edges between binding site residues (B–B) and an 8 Å cutoff for all other edges (B–NB and NB–NB). In these hybrid 24 Å | 8 Å-cutoff graphs, only the number of B–B edges increases compared to the standard 8 Å-cutoff graphs. The GNN trained on the human hybrid graphs (24 Å | 8 Å GNN) substantially reduced the MCC loss compared with the 8 Å GNN (**Fig. S2**). The areas under the ROC and precision-recall curves also increase, reaching at least 0.95 and 0.86, respectively, for the rat protein graphs (**Fig. 3b-c**).

**Fig. 3:**
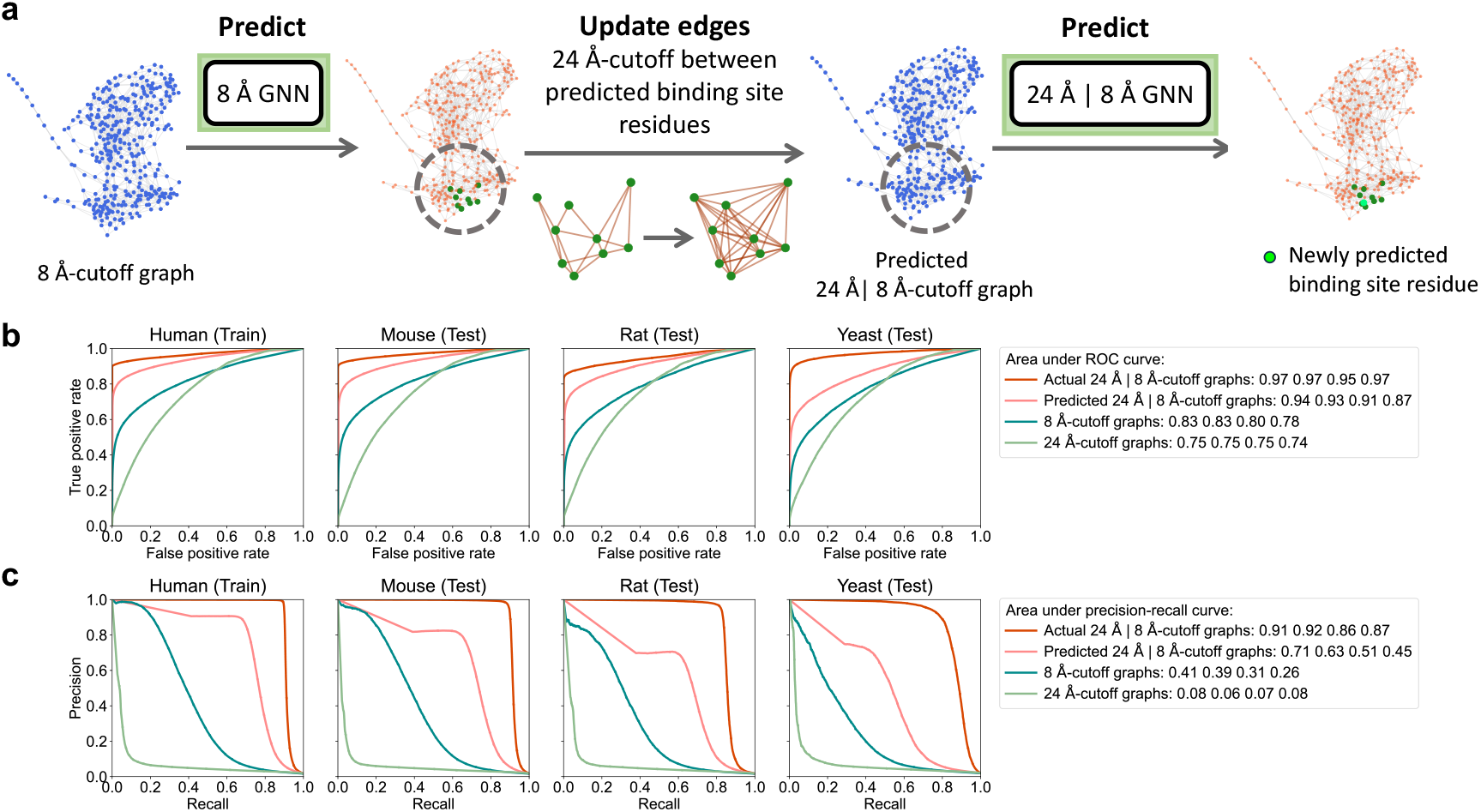
24 Å | 8 Å GNN performs better on predicted 24 Å | 8 Å-cutoff graphs than on the standard 8 Å- or 24 Å-cutoff graphs. **a** Procedures of generating predicted 24 Å | 8 Å-cutoff cutoff graphs and testing 24 Å | 8 Å GNN on these predicted hybrid cutoff graphs. The GNN trained on 8 Å-cutoff human protein graphs (8 Å GNN) is applied to predict potential binding site residues, between which the edges are then enriched using a 24 Å cutoff to generate predicted hybrid cutoff graphs. **b** Receiver operating characteristic (ROC) curves and **c** precision-recall curves for 24 Å | 8 Å GNN tested on actual hybrid cutoff graphs (red), predicted hybrid cutoff graphs (orange), 8 Å-cutoff graphs (green), and 24 Å-cutoff graphs (light green).

As a control, we also construct inverted hybrid 8 Å | 24 Å-cutoff graphs, where B–B edges are sparse but the other edges are dense. This design does not improve the GNN’s performance. The MCC loss and the areas under the ROC and precision-recall curves are comparable to those of 24 Å GNN (**Fig. S3**). Together, these results indicate that performance gains are not due to simply adding edges but specifically to increasing connectivity between binding site residues.

Finally, we examine whether 24 Å | 8 Å GNN generalizes to other cutoff graphs. Considering that binding site residues are the prediction targets, it is impractical to generate graphs with more edges between binding site residues beforehand. Therefore, we generate predicted hybrid 24 Å | 8 Å-cutoff graphs by using 8 Å GNN to predict binding site residues from 8 Å-cutoff graphs. We then updated edges using 24 Å cutoff for the distances between the predicted binding site residues while keeping other edges unchanged (**Fig. 3a**). Although errors in residue classification introduce noise and reduce performance relative to the actual hybrid cutoff graphs, the predicted hybrid cutoff graphs still outperform both standard 8 Å- and 24 Å-cutoff graphs (**Fig. 3b-c**). This result demonstrates the potential for enhancing GNN’s performance by iteratively increasing connectivity between accurately predicted binding site residues.

### Sequence-based features and structure-based features contribute both overlapping and distinct information for binding site residue prediction

The GNNs evaluated until this point are performed on graphs with nodes characterized by both sequence-based and structure-based features. Considering that sequence-based features, the learned representations of the ESM-2 language model, can capture protein secondary and tertiary structures [33, 34], it is unclear whether structure-based features, the backbone dihedral angles *ϕ* and *ψ*, provide additional information for binding site residue prediction. To disentangle the impacts of these two types of features on binding site residue prediction, we retrain 8 Å GNN on the human protein graphs using either only sequence-based node features or only structure-based node features. 8 Å GNN is trained on the human protein graphs using both sequence-based and structure-based features. We name the two retrained GNNs ‘sequence-based only GNN’ and ‘structure-based only GNN’, respectively. The MCC loss of sequence-based only GNN is marginally lower than the loss of 8 Å GNN. On the other hand, the MCC loss of structure-based only GNN barely decreases (**Fig. S4a**). To investigate whether the hyperparameters of 8 Å GNN limit its ability to learn from either type of features, we reoptimize the hyperparameters for the two GNNs (**Table S2**). However, the loss trends are largely unaffected (**Fig. S4b**).

Despite the poor MCC loss, the area under the ROC curve of structure-based only GNN is substantially greater than random chance at 0.65 (**Fig. 4a**), indicating that the GNN has some ability to distinguish between binding and non-binding site residues. On the other hand, the area under the precision-recall curve is only slightly higher than random at 0.03 (**Fig. 4b**). The precision is low when the recall is high, indicating a strong bias of structure-based only GNN toward false positive predictions. To explore further, we compare the true positives and false positives predicted by structure-based only GNN and by sequence-based only GNN using a threshold of 0.5 (**Fig. 4c**). Apart from the numerous false positives, structure-based only GNN also makes a substantial number of true positive predictions where sequence-based only GNN fails, especially for the binding site residues of the yeast proteins. This result demonstrates that there are some unique contributions from the backbone dihedral angles to binding that is not captured by the ESM-2 residue representations.

**Fig. 4:**
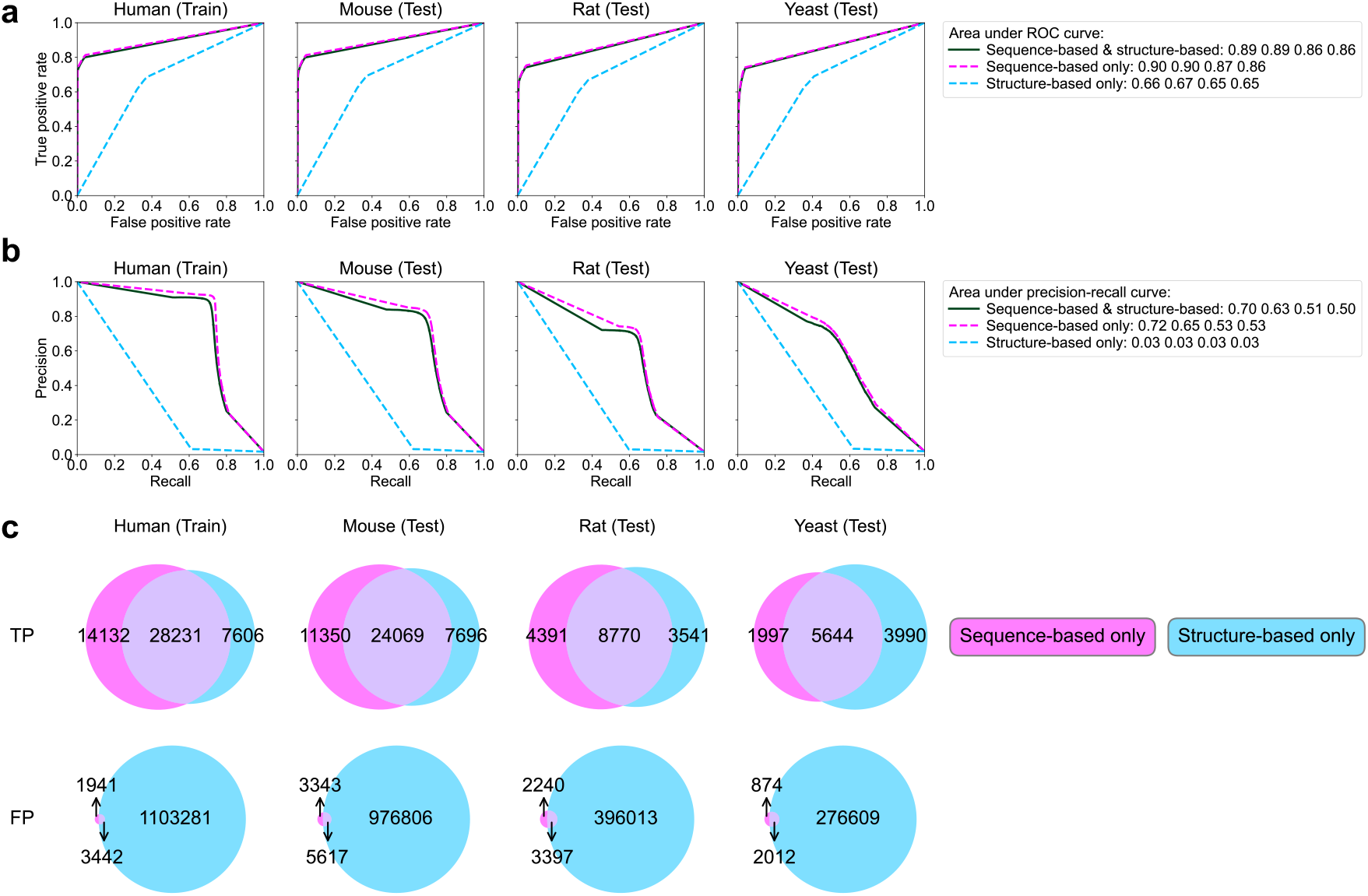
Structure-based features using protein backbone dihedral angles *ϕ* and *ψ* exhibit bias toward positive predictions. **a** Receiver operating characteristic (ROC) curves and **b** precision-recall curves for three GNNs after the last training epoch. The first GNN is the reference 8 Å GNN, which is trained and tested on graphs with nodes characterized by both sequence-based and structure-based features (dark green solid line). The second and third GNNs are 8 Å GNN retrained and tested on graphs using either sequence-based features only (magenta dashed line) or structure-based features only (cyan dashed line), respectively. **c** Venn diagrams comparing true positives (TP) and false positives (FP) predicted by sequence-based only GNN and structure-based only GNN in each dataset using a threshold of 0.5.

## Discussion

Binding site residues are essential for protein function but difficult to predict due to their low occurrence and divergent evolution of protein sequence and structure. Despite this complexity, protein sequence and structure remain informative resources to identify binding site residues. The challenge then lies in optimizing sequence-based and structure-based features that determine binding. In this study, we investigate the effect of different sequence-based and structure-based features on binding site residue prediction with graph neural networks (GNNs). We derive sequence-based and structure-based features from inter-residue distances, residue representations, and residue backbone geometry.

Dense connectivity between binding site residues and sparse connectivity to non-binding site residues enhances binding site residue prediction. Using different cutoffs on inter-residue distances to construct edges, we determine that 8 Å cutoff is most effective to describe pairwise residue interactions for binding. We find that increasing cutoff distances introduces disproportionately more edges to non-binding site residues and reduces the GNN’s performance. Conversely, hybrid cutoffs that selectively increase connectivity between true binding site residues substantially enhance the GNN’s performance, with the largest improvement in the yeast proteins where the area under precision-recall curve increases from 0.50 to 0.87. This finding can guide future studies on methods to iteratively update edges and optimize the precision of binding site residue prediction. The effectiveness of such methods depends on minimizing false positives to prevent erroneously adding more edges to non-binding site residues. Moreover, inter-residue distances may be represented as edge features to enrich description of pairwise residue interactions beyond discrete cutoffs.

ESM-2 residue representations are informative for binding site residue prediction, while residue backbone geometry provides minimal complement to the GNN’s performance. We evaluate feature contributions from ESM-2 residue representations and residue backbone geometry. The residue representations are the information that a pre-trained ESM-2 model learned during training on millions of protein sequences from UniProtKB [34]. As these sequence-derived representations may implicitly carry some structural information [24, 33, 35], we consider residue backbone geometry as a complementary structure-derived representation that minimizes sequence contributions by residue sidechains. We find that using the residue representations alone, the GNN’s performance is as good as using both residue representations and backbone geometry. In contrast, residue backbone geometry leads to overprediction of binding site residues, indicating that having a similar geometry of a binding site without direct sidechain information is insufficient for accurate predictions. Furthermore, reliability of backbone geometry closely depends on the accuracy of the AlphaFold predicted structures. Incorporating AlphaFold’s local confidence score into structure-based features may calibrate noise from the uncertainty of predicted protein structures. These results motivate further feature engineering that seeks for more meaningful structure-derived representations to complement ESM-2 residue representations.

In this study, all GNNs are trained on the human protein graphs, yet they capture binding site signals of the mouse, rat, and even yeast proteins. For instance, the area under precision-recall curve drops by only 0.20 from human (0.70) to yeast (0.50) using 8 Å cutoff. Comparative genome analysis estimated that 80% to 90% of the mouse and rat genes have orthologs in the human genome [40, 41]. Although only about one-third of the yeast genes have orthologs in the human genome, nearly half of the yeast genes required for yeast cell growth can be functionally replaced by their human orthologs [42]. Given that binding is an essential mechanism of protein function, it is likely that binding sites are retained in some of these genes, enabling the GNN’s generalizability. Our study not only demonstrates the relevant information derived from protein sequences and structures on binding site residue prediction but also provides evidence for evolutionary conservation of binding site residues.

## Methods

### Constructing training and test datasets

Binding site residue annotations and the associated protein accession numbers of four model organism proteomes are collected from the genome annotation tracks of Universal Protein Resource Knowledgebase (UniProtKB) Release 2024_05 [23]. The four organisms are *Homo sapiens* (human), *Mus musculus* (mouse), *Rattus norvegicus* (rat), and *Saccharomyces cerevisiae* (yeast). The three-dimensional (3D) structures, including sequences and Cartesian coordinates, of the four organism proteomoes are retrieved from the corresponding proteome archive files of the AlphaFold Protein Structure Database (AlphaFold DB version 4) [24, 25]. Structures of UniProtKB entries that are missing from the AlphaFold archive files are retrieved individually from the AlphaFold DB version 4 website using protein accession numbers. UniProtKB entries with a sequence length greater than 2,700 residues are excluded. Each UniProtKB entry kept is represented as an undirected graph using PyTorch Geometric [43]. The number of protein graphs in each dataset is 5994 (human), 5223 (mouse), 2177 (rat), and 1362 (yeast). The human dataset is used for hyperparameter optimization and training, while the other three datasets are used for testing.

### Representing proteins as graphs

In mathematical terms, a graph *G* is an ordered pair

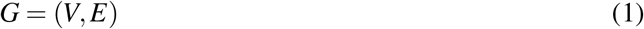

where *V* = {*v*_1_,…, *v*_*n*_} is a set of *n* nodes and *E* ⊂ *V* × *V* is the set of all edges that connect two nodes. Each edge is an unordered pair (*v*_*i*_, *v* _*j*_) *E*. Associated to each node *v*_*i*_ is a *p*-dimensional feature vector **x**_*i*_ ∈ ℝ^*p*^. The topology of a graph is described through an adjacency matrix *A*, an *n × n* square matrix whose entries are associated with edges of the graph according to the following rule:

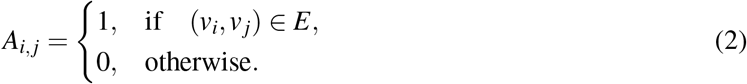

Two nodes *v*_*i*_ and *v* _*j*_ are called neighbors if and only if *A*_*i, j*_ = 1, that is, if and only if (*v*_*i*_, *v* _*j*_) ∈ *E*. For each *i* ∈ {1,…, *n*}, the neighborhood *N*_*i*_ of *v*_*i*_ is the set of all nodes that share an edge with *v*_*i*_:

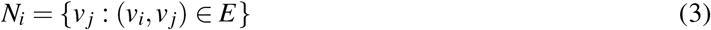

Every protein in the four datasets above is represented as a graph, with each node representing a protein residue, each edge representing interaction between a pair of residues, and the number of nodes *n* being equal to the protein sequence length. Each node is equipped with an input feature vector and a binary label

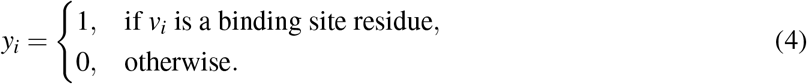

A node pair (*v*_*i*_, *v*_*j*_) is assigned an edge if the distance between C_*β*_ atoms of the two residues (C_*α*_ for glycine) is less than an arbitrary cutoff distance *d* > 0. We consider edge cutoffs of 8 Å, 12 Å, 16 Å, 20 Å, and 24 Å.

Three input feature vectors are considered: (i) a vector with only sequence-based features, (ii) a vector with only structure-based features, and (iii) a vector with both sequence-based and structure-based features. To construct sequence-based features, we apply the pre-trained ESM-2 masked protein language model with 33 layers and 650 million parameters (esm2_t33_650M_UR50D) [33] to each protein sequence and generate a 1,280-dimensional embedding vector for each residue. To construct structure-based features, we calculate the backbone dihedral angles *ϕ* and *ψ*, a two-dimensional vector per residue, from the protein Cartesian coordinates using MDAnalysis [44]. The dihedral angles of the first and last residues are duplicated from the dihedral angles of their immediately adjacent residues, i.e., the second residue and the second to the last residue, respectively.

### Graph neural network architecture

The graph neural network (GNN) architecture used in this work is shown in **Fig. 1a**. The first layer in this architecture is the input layer, which contains the input features of a protein. For each node *v*_*i*_, these features are collected into an input feature vector 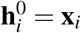. Then, for each *k* ∈ {1,…, *K*}, the *k*^th^ hidden layer consists of (1) a graph convolutional operator for sharing information between nodes, (2) a ReLU activation function, which introduces nonlinearity, and (3) a dropout operator that randomly removes a fraction of nodes during training to prevent overfitting. More specifically, the steps to get from 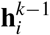 to 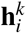 are

1a. Aggregate information from neighbors: For each *i* ∈*{*1,…, *n}*, the node *v*_*i*_ collects the embedded features of its neighbors via an aggregation function:

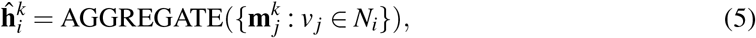

where 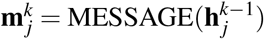 is a message obtained from the neighboring node *v* _*j*_ ∈ *N*_*i*_. The vector 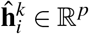is the embedded hidden feature vector of node *v*_*i*_ in the *k*^th^ hidden layer.

1b. Update hidden state information: With 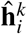collected, the nodal feature of node *v*_*i*_ is updated:

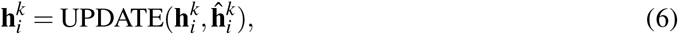

where UPDATE is a differentiable function that combines aggregated messages 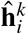collected from the neighborhood *N*_*i*_ with the feature vector 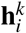of the current layer.

2. Apply the ReLU activation:

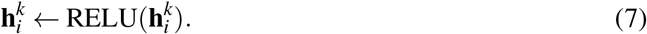

3. Apply the dropout regularization on a fraction of nodes in the hidden layer during training:

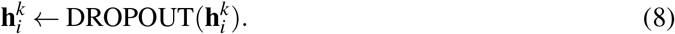

Steps (1a) and (1b) above together form the graph convolutional operator. Their specific implementation is based on the GCNConv operator described in [36]. The final output layer consists of another GCNConv operator followed by a sigmoid activation function that generates the probability *p*_*i*_ that residue *i* is a binding site residue. We implement the GNN architecture in PyTorch Geometric [43].

### Loss function

We use the loss function [37]

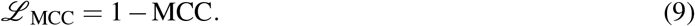

to measure errors between predicted probabilities and actual labels. Here MCC is the Matthews Correlation Coefficient [45]

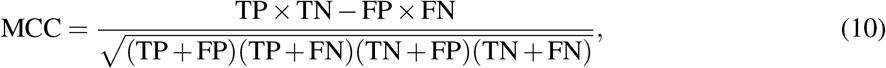

where the quantities

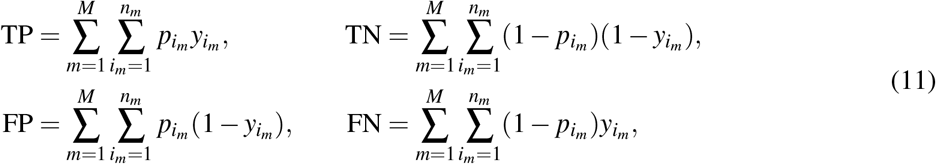

are computed over a dataset with *M* proteins labeled by *m* ∈ {1,…, *M*}. Each protein has *n*_*m*_ residues and for each *i*_*m*_ ∈ {1,…, *n*_*m*_}, 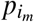is the predicted probability that residue *i*_*m*_ of protein *m* is a binding site residue. Meanwhile, 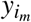is the actual residue label. The formulas in (11) ensure that the *ℒ* _MCC_ is differentiable with respect to the predicted probabilities 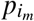, which is a requirement for training. If *p*_*i*_ ∈ {0, 1} for all *i* ∈{1,…, *n*}, these formulas recover the usual values for the numbers of true positives, true negatives, false positives, and false negatives.

### GNN hyperparameter optimization and training

We perform hyperparameter optimization using Optuna [46] to maximize the MCC. For each choice of cutoff distance and input features (sequence-based, structure-based, or both), we consider a search space with five hyperparameters: (i) the number of hidden layers, (ii) the number of neurons in each hidden layer, (iii) the dropout rate in each hidden layer, (iv) the type of optimizer, and (v) the learning rate.

The initial search space spans (i) one to eight hidden layers, (ii) four to 128 neurons per layer, (iii) a dropout rate between 0.2 and 0.8 per layer, (iv) an Adam [47] or RMSprop [48] optimizer, and (v) a learning rate between 10^-5^ and 10^-1^. The hyperparameter optimization for the choices of 8 Å- and 12 Å-cutoffs result in GNNs with one hidden layer of 128 neurons. Because the setting of 128 neurons is the upper bound of the search space, we initiate another round of optimization just for these two cutoff distances, with an updated search space having (i) one to two hidden layers and (ii) 128 to 512 neurons per layer, while keeping the ranges of the other hyperparameters the same.

The results in **Table S1** are for the input feature vector that includes both sequence-based and structure-based features, and the optimization is performed independently for each cutoff distance. The results in **Table S2** are for a fixed cutoff of 8 Å, and the optimization is performed independently for either sequence-based features only or structure-based features only.

All optimizations are performed using the human dataset with 5-fold cross validation for 100 trials. Each fold is trained for 100 epochs using a batch size of 128. MCC values are averaged over the five folds for each trial, and then the maximum of these averages over 100 such trials is recorded. To mitigate against outliers, the optimization is performed three times for each choice of cutoff distance and input features. While each instance of the optimization might yield slightly different hyperparameters and maximum MCC, the deviation in the maximum MCC is relatively small. See **Tables S1-S2** for the average and standard deviation of the maximum MCC over the three instances.

For each choice of cutoff distance and input features, the hyperparameters that produce the maximum MCC across all three instances of the optimization are used for subsequent training and testing. The associated GNNs are retrained on the entire human dataset for 500 epochs using a batch size of 128. The model parameters are saved after every epoch. To ensure effective training and prevent overfitting, the GNNs are evaluated at each epoch on all four datasets using the MCC loss in formula (9).

### Metrics

The GNNs after the last training epoch are evaluated on all four datasets using receiver operating characteristic (ROC) curves and precision-recall curves implemented in scikit-learn [49]. An ROC curve is a plot of the true positive rate (TPR) against the false positive rate (FPR) for each value of a varying threshold *τ* ∈ [0, 1], while a precision-recall curve is a plot of the precision against recall (TPR) for each value of *τ*. The relevant formulas are

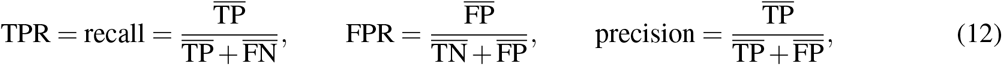

where, similar to (11),

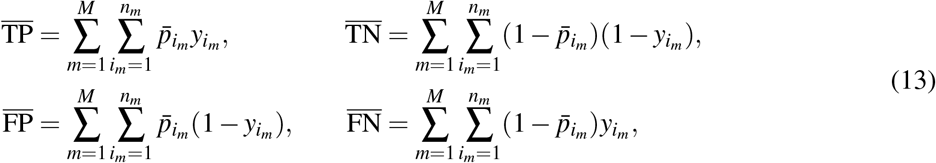

and *y*_*i*_ is the actual label at residue *i*. However, 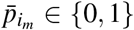 is now a rounded value of the prediction probability that depends on *τ*:

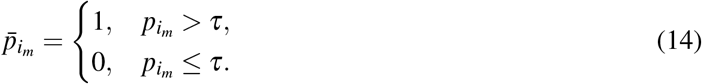

For a given prediction 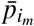, 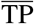is the number of correctly classified binding site residues, 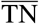 is the number of correctly classified non-binding site residues, 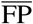 is the number of incorrectly classified binding site residues, and 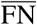 is the number of incorrectly classified non-binding site residues.

### Comparison with random guessing

We consider two random guessing strategies. One strategy, named “random (50%)”, assumes each residue has a 50% chance of being a binding site residue. The other strategy, named “random (1.6% – 1.9%)”, uses a prior knowledge of the percentage of binding site residues in each dataset and randomly selected 1.7%, 1.6%, 1.7%, and 1.9% of the residues in each protein graph as binding site residues for the human, mouse, rat, and yeast datasets, respectively. Both strategies are applied to each of the four datasets. The same metrics, MCC loss, ROC curve, and precision-recall curve, are computed as in testing.

### Hardware resources

Hyperparameter optimization is performed using six Intel Xeon vCPUs and one Nvidia Tesla V100 GPU with 32 GB memory. Training and testing are performed on a 14-core Apple M3 Max laptop with 36 GB memory.

## Data availability

The four datasets and GNNs developed in this work are available at https://github.com/serenachen0/PBSGNN.

## Code availability

All the codes developed in this work are available at https://github.com/serenachen0/PBSGNN.

## Acknowledgments

S.H.C. thanks David R. Bell (FNLCR) for fruitful discussion and comments on the manuscript, Chris Layton (ORNL), Daniel Dewey (ORNL), and Daniel Collins (ORNL) for technical support on the ORNL Research Cloud. The authors also thank Thomas Strohmer (UC Davis), Jonathan Forstater (UC Davis), Bao Wang (Univ. Utah), Justin M. Baker (Univ. Utah), and Shih-Hsin Wang (Univ. Utah) for helpful discussion about this work.

This work is supported by the Office of Advanced Scientific Computing Research of the U.S. Department of Energy under contract No. DE-SC0023490. This work uses resources from the ORNL Research Cloud Infrastructure at the Oak Ridge National Laboratory, which is supported by the Office of Science of the U.S. Department of Energy under Contract No. DE-AC05-00OR22725.

## Author contributions

S.H.C: conceptualization (lead), data curation (lead), formal analysis (lead), funding acquisition (supporting), investigation (lead), methodology (lead), project administration (supporting), resources (lead), software (lead), validation (lead), visualization (lead), writing - original draft preparation (lead), writing - review and editing (equal). M.L.P: conceptualization (supporting), funding acquisition (supporting), methodology (supporting), project administration (supporting), writing - review and editing (equal). C.D.H: conceptualization (supporting), funding acquisition (lead), project administration (lead), writing - review and editing (equal).

## References

[1] Xia, Y., Xia, C.-Q., Pan, X. & Shen, H.-B. Graphbind: protein structural context embedded rules learned by hierarchical graph neural networks for recognizing nucleic-acid-binding residues. Nucleic acids research 49, e51–e51 (2021).

[2] Yuan, Q., Chen, J., Zhao, H., Zhou, Y. & Yang, Y. Structure-aware protein–protein interaction site prediction using deep graph convolutional network. Bioinformatics 38, 125–132 (2022).

[3] Wu, H. et al. Spatom: a graph neural network for structure-based protein–protein interaction site prediction. Briefings in Bioinformatics 24, bbad345 (2023).

[4] Han, J. et al. Geonet enables the accurate prediction of protein-ligand binding sites through interpretable geometric deep learning. Structure 32, 2435–2448 (2024).

[5] Yuan, Q., Tian, C. & Yang, Y. Genome-scale annotation of protein binding sites via language model and geometric deep learning. Elife 13, RP93695 (2024).

[6] Piana, S. et al. Evaluating the effects of cutoffs and treatment of long-range electrostatics in protein folding simulations. PLoS One 7, e39918 (2012).

[7] Salamanca Viloria, J., Allega, M. F., Lambrughi, M. & Papaleo, E. An optimal distance cutoff for contact-based protein structure networks using side-chain centers of mass. Scientific reports 7, 2838 (2017).

[8] Jiang, M. et al. Drug–target affinity prediction using graph neural network and contact maps. RSC advances 10, 20701–20712 (2020).

[9] Zhang, S. et al. Ss-gnn: a simple-structured graph neural network for affinity prediction. ACS omega 8, 22496–22507 (2023).

[10] Airas, J. & Zhang, B. Scaling graph neural networks to large proteins. Journal of chemical theory and computation 21, 2055–2066 (2025).

[11] Lichtarge, O., Bourne, H. R. & Cohen, F. E. An evolutionary trace method defines binding surfaces common to protein families. Journal of molecular biology 257, 342–358 (1996).

[12] Illergård, K., Ardell, D. H. & Elofsson, A. Structure is three to ten times more conserved than sequence—a study of structural response in protein cores. Proteins: Structure, Function, and Bioinformatics 77, 499–508 (2009).

[13] Stern, D. L. The genetic causes of convergent evolution. Nature Reviews Genetics 14, 751–764 (2013).

[14] Konc, J. & Janežič, D. Probis algorithm for detection of structurally similar protein binding sites by local structural alignment. Bioinformatics 26, 1160–1168 (2010).

[15] Tseng, Y. Y. & Li, W.-H. Evolutionary approach to predicting the binding site residues of a protein from its primary sequence. Proceedings of the National Academy of Sciences 108, 5313–5318 (2011).

[16] Jiménez, J., Doerr, S., Martínez-Rosell, G., Rose, A. S. & De Fabritiis, G. Deepsite: protein-binding site predictor using 3d-convolutional neural networks. Bioinformatics 33, 3036–3042 (2017).

[17] Mahmud, M., Kaiser, M. S., McGinnity, T. M. & Hussain, A. Deep learning in mining biological data. Cognitive computation 13, 1–33 (2021).

[18] Islam, M. T. et al. Revealing hidden patterns in deep neural network feature space continuum via manifold learning. Nature Communications 14, 8506 (2023).

[19] Gligorijević, V. et al. Structure-based protein function prediction using graph convolutional networks. Nature communications 12, 3168 (2021).

[20] NIAID Visual & Medical Arts. (10/7/2024). Human Female Outline_type VI. NIAID NIH BIOART Source.bioart.niaid.nih.gov/bioart/227.

[21] NIAID Visual & Medical Arts. (10/7/2024). Lab Mouse. NIAID NIH BIOART Source. bioart.niaid.nih.gov/bioart/281.

[22] NIAID Visual & Medical Arts. (10/7/2024). Black Rat. NIAID NIH BIOART Source. bioart.niaid.nih.gov/bioart/54.

[23] The UniProt Consortium. Uniprot: the universal protein knowledgebase in 2025. Nucleic Acids Research 53, D609–D617 (2024).

[24] Jumper, J. et al. Highly accurate protein structure prediction with alphafold. nature 596, 583–589 (2021).

[25] Varadi, M. et al. Alphafold protein structure database in 2024: providing structure coverage for over 214 million protein sequences. Nucleic acids research 52, D368–D375 (2024).

[26] Poux, S. et al. On expert curation and scalability: Uniprotkb/swiss-prot as a case study. Bioinformatics 33, 3454–3460 (2017).

[27] Blum, M. et al. Interpro: the protein sequence classification resource in 2025. Nucleic Acids Research 53, D444–D456 (2025).

[28] Fasoulis, R., Paliouras, G. & Kavraki, L. E. Graph representation learning for structural proteomics. Emerging Topics in Life Sciences 5, 789–802 (2021).

[29] Maier, J. A. et al. ff14sb: improving the accuracy of protein side chain and backbone parameters from ff99sb. Journal of chemical theory and computation 11, 3696–3713 (2015).

[30] Huang, J. et al. Charmm36m: an improved force field for folded and intrinsically disordered proteins. Nature methods 14, 71–73 (2017).

[31] Chen, S. H. & Bell, D. R. Evolution of thyroglobulin loop kinetics in epcam. Life 11, 915 (2021).

[32] Chen, S. H., Weiss, K. L., Stanley, C. & Bhowmik, D. Structural characterization of an intrinsically disordered protein complex using integrated small-angle neutron scattering and computing. Protein Science 32, e4772 (2023).

[33] Lin, Z. et al. Evolutionary-scale prediction of atomic-level protein structure with a language model. Science 379, 1123–1130 (2023).

[34] Rives, A. et al. Biological structure and function emerge from scaling unsupervised learning to 250 million protein sequences. Proceedings of the National Academy of Sciences 118, e2016239118 (2021).

[35] Baek, M. et al. Accurate prediction of protein structures and interactions using a three-track neural network. Science 373, 871–876 (2021).

[36] Kipf, T. N. & Welling, M. Semi-supervised classification with graph convolutional networks. In International Conference on Learning Representations (2017). URL https://openreview.net/forum?id=SJU4ayYgl.

[37] Abhishek, K. & Hamarneh, G. Matthews correlation coefficient loss for deep convolutional networks: Application to skin lesion segmentation. In 2021 IEEE 18th International Symposium on Biomedical Imaging (ISBI), 225–229 (IEEE, 2021).

[38] Chicco, D. & Jurman, G. The advantages of the matthews correlation coefficient (mcc) over f1 score and accuracy in binary classification evaluation. BMC genomics 21, 6 (2020).

[39] Chicco, D., Tötsch, N. & Jurman, G. The matthews correlation coefficient (mcc) is more reliable than balanced accuracy, bookmaker informedness, and markedness in two-class confusion matrix evaluation. BioData mining 14, 13 (2021).

[40] Mouse Genome Sequencing Consortium. Initial sequencing and comparative analysis of the mouse genome. Nature 420, 520–562 (2002).

[41] Mullins, L. J. & Mullins, J. J. Insights from the rat genome sequence. Genome biology 5, 1–3 (2004).

[42] Kachroo, A. H. et al. Systematic humanization of yeast genes reveals conserved functions and genetic modularity. Science 348, 921–925 (2015).

[43] Fey, M. & Lenssen, J. Fast graph representation learning with pytorch geometric. In ICLR 2019 Workshop on Representation Learning on Graphs and Manifolds (2019).

[44] Michaud-Agrawal, N., Denning, E. J., Woolf, T. B. & Beckstein, O. Mdanalysis: a toolkit for the analysis of molecular dynamics simulations. Journal of computational chemistry 32, 2319–2327 (2011).

[45] Matthews, B. W. Comparison of the predicted and observed secondary structure of t4 phage lysozyme. Biochimica et Biophysica Acta (BBA)-Protein Structure 405, 442–451 (1975).

[46] Akiba, T., Sano, S., Yanase, T., Ohta, T. & Koyama, M. Optuna: A next-generation hyperparameter optimization framework. In Proceedings of the 25th ACM SIGKDD International Conference on Knowledge Discovery and Data Mining (2019).

[47] Adam, K. D. B. J. et al. A method for stochastic optimization. arXiv preprint 1412.6980 1412 (2014).

[48] Shi, N. & Li, D. RMSprop converges with proper hyperparameter. In International conference on learning representation (2021). URL https://openreview.net/forum?id=3UDSdyIcBDA.

[49] Pedregosa, F. et al. Scikit-learn: Machine learning in Python. Journal of Machine Learning Research 12, 2825–2830 (2011).

