## Supplementary Information for "Enhancing Protein Binding Site Residue Prediction with Graph Neural Networks: Impacts of Cutoff Distance and Feature Selection"

**Table S1: Optimal hyperparameters of graph neural networks for protein graphs constructed by different cutoff distances.**

| Optimization #1 | | | | | | | | |
|---|---|---|---|---|---|---|---|---|
| **Hyperparameter** | **Search space** | **8 Å** | **12 Å** | **16 Å** | **20 Å** | **24 Å** | **24 Å ǀ 8 Å‡** | **8 Å ǀ 24 Å‡** |
| Number of hidden layers | 1 – 8 | 1 | 1 | 1 | 1 | 1 | 1 | 1 |
| Number of neurons in each hidden layer | 4 – 128 | 128 | 128 | 59 | 44 | 9 | 98 | 12 |
| Dropout rate in each hidden layer | 0.2 – 0.8 | 0.586 | 0.292 | 0.263 | 0.693 | 0.507 | 0.357 | 0.260 |
| Optimizer | Adam, RMSprop | Adam | Adam | Adam | Adam | Adam | Adam | Adam |
| Learning rate | $10^{-5}$ – $10^{-1}$ | 0.007 | 0.022 | 0.027 | 0.014 | 0.030 | 0.019 | 0.027 |
| Maximum MCC | | 0.594 | 0.519 | 0.447 | 0.391 | 0.342 | 0.824 | 0.330 |
| Average of maximum MCC (SD) | | 0.592 (0.001) | 0.516 (0.003) | 0.445 (0.002) | 0.384 (0.006) | 0.329 (0.014) | 0.812 (0.011) | 0.316 (0.012) |

| Optimization #2 | | | |
|---|---|---|---|
| **Hyperparameter** | **Search space** | **8 Å** | **12 Å** |
| Number of hidden layers | 1 – 2 | 1 | 1 |
| Number of neurons in each hidden layer | 128 – 512 | 297 | 322 |
| Dropout rate in each hidden layer | 0.2 – 0.8 | 0.439 | 0.295 |
| Optimizer | Adam, RMSprop | Adam | Adam |
| Learning rate | $10^{-5}$ – $10^{-1}$ | 0.010 | 0.012 |
| Maximum MCC | | 0.602 | 0.522 |
| Average of maximum MCC (SD) | | 0.596 (0.008) | 0.520 (0.002) |

*All graph nodes are characterized by both sequence-based and structure-based features. For each evaluated cutoff distance, three replicas of 100-trial hyperparameter search are performed, and the average and standard deviation of maximum MCC are reported. The optimal hyperparameter set and the associated MCC are also reported.

‡x Å ǀ y Å denotes using x Å cutoff to define edges between binding site residues and y Å cutoff to define edges between binding and non-binding site residues as well as between non-binding site residues.

**Table S2: Optimized hyperparameters of graph neural networks for protein graphs characterized by either only sequence-based node features or only structure-based node features.**

| Hyperparameter | Search space | Sequence-based only | Structure-based only |
|---|---|---|---|
| Number of hidden layers | 1 – 8 | 1 | 4 |
| Number of neurons in each hidden layer | 4 – 128 | 113 | 124, 113, 8, 102 |
| Dropout rate in each hidden layer | 0.2 – 0.8 | 0.618 | 0.234, 0.349, 0.406, 0.356 |
| Optimizer | Adam, RMSprop | Adam | RMSprop |
| Learning rate | $10^{-5}$ – $10^{-1}$ | 0.017 | 0.001 |
| Maximum MCC | | 0.609 | 0.098 |
| Average of maximum MCC (SD) | | 0.607 (0.001) | 0.095 (0.006) |

*Edges are defined using 8 Å cutoff. Three replicas of 100-trial hyperparameter search are performed for each set of protein graphs, and the average and standard deviation of maximum MCC are reported. The optimal hyperparameter set and the associated MCC are also reported.

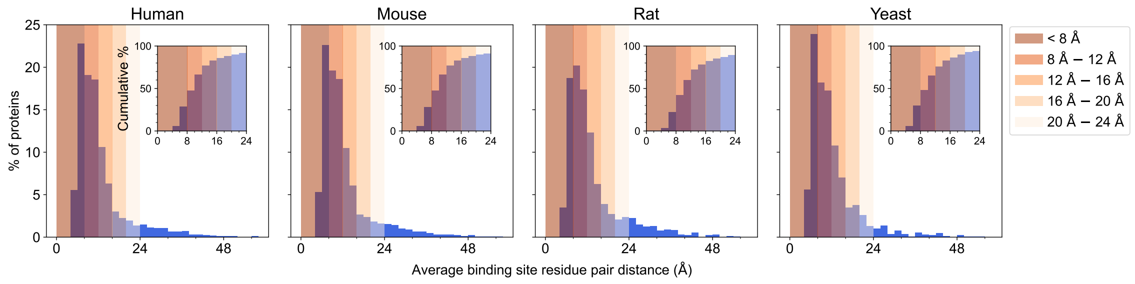


**Fig. S1: Average distance between binding site residue pairs levels off by 24 Å for all proteins in each of the four model organism datasets.** Histograms illustrate the distribution of proteins in the human, mouse, rat, and yeast datasets across average binding site residue pair distances ranging from 0 Å to 60 Å at an interval of 2 Å. The insets provide the cumulative protein percentages for the distance up to 24 Å. To assist visualization, shaded bars with a decreasing color gradient are included for the distances less than 8 Å (< 8 Å) and between 8 Å and 24 Å at an interval of 4 Å (8 Å – 12 Å, 12 Å – 16 Å, 16 Å – 20 Å, 20 Å – 24 Å, respectively).

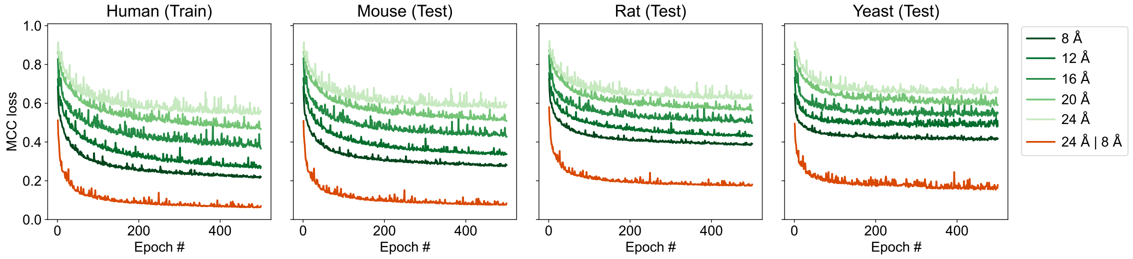


**Fig. S2: Loss curve increases with increasing cutoff distances and decreases with more edges between binding site residues.** Six graph neural networks (GNNs) are evaluated by Matthews Correlation Coefficient (MCC) loss over 500 epochs. Each GNN is trained on the human protein graphs and tested on the mouse, rat, and yeast protein graphs constructed with either one or two cutoff distances throughout the proteins. For graphs of one cutoff distance, the cutoffs are 8 Å, 12 Å, 16 Å, 20 Å, and 24 Å, and their loss curves are shown as a series of green shades from dark to light. For graphs of two cutoff distances, the cutoffs are 24 Å between binding site residues and 8 Å between binding and non-binding site residues as well as between non-binding site residues (24 Å | 8 Å, red).

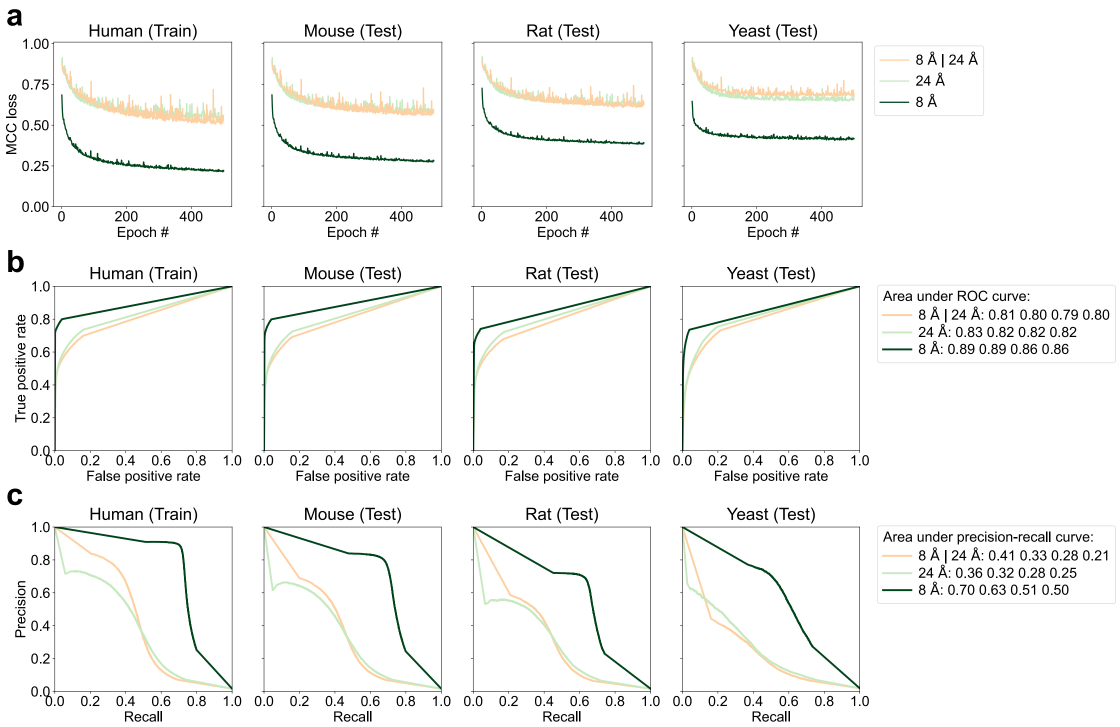


**Fig. S3: Adding more edges to non-binding site residues decreases the performance of graph neural networks.** **a** MCC loss curve, **b** ROC curve, and **c** precision-recall curve for the GNN trained and tested on the graphs constructed by 8 Å cutoff between binding site residues and 24 Å cutoff between binding and non-binding site residues as well as between non-binding site residues (8 Å | 24 Å, beige). Performance of the GNNs trained and tested on the graphs constructed by 8 Å cutoff (dark green) or 24 Å cutoff (light green) throughout the proteins are included for reference.

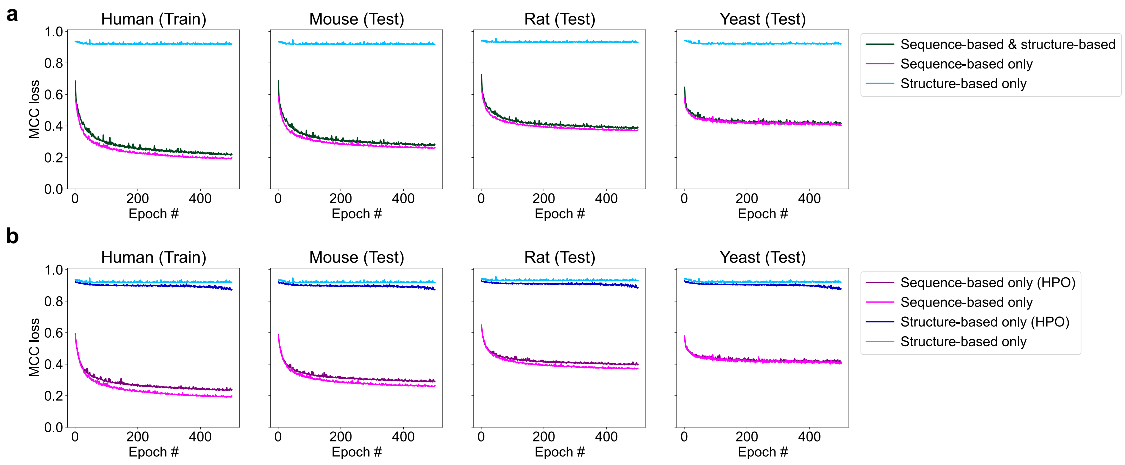


**Fig. S4: Structure-based features based on protein backbone dihedral angles $\phi$ and $\psi$ have a minimal impact on the performance of graph neural networks.** Performance is measured by Matthews Correlation Coefficient (MCC) loss of a graph neural network (GNN) during training on the human dataset and testing on the mouse, rat, and yeast datasets over 500 epochs. **a** MCC loss curves of three GNNs trained and tested on the 8 Å-cutoff graphs with nodes characterized by both sequence-based and structure-based features (dark green, same as the 8 Å-cutoff loss curve in Fig. S2), by sequence-based features only (magenta), and by structure-based features only (cyan). All three GNNs have the same hyperparameters of the GNN optimized with both sequence-based and structure-based node features and 8 Å cutoff. **b** MCC loss curves of the hyperparameter-optimized (HPO) GNNs trained and tested on the 8 Å-cutoff graphs with nodes characterized by either sequence-based features only (dark purple) or by structure-based features only (dark blue). The loss curves of the GNNs before hyperparameter optimization (magenta and cyan) are the same as in **a** and included here for reference.
